# Interactions between species introduce spurious associations in microbiome studies

**DOI:** 10.1101/176677

**Authors:** Rajita Menon, Vivek Ramanan, Kirill S. Korolev

## Abstract

Microbiota contribute to many dimensions of host phenotype, including disease. To link specific microbes to specific phenotypes, microbiome-wide association studies compare microbial abundances between two groups of samples. Abundance differences, however, reflect not only direct associations with the phenotype, but also indirect effects due to microbial interactions. We found that microbial interactions could easily generate a large number of spurious associations that provide no mechanistic insight. Using techniques from statistical physics, we developed a method to remove indirect associations and applied it to the largest dataset on pediatric inflammatory bowel disease. Our method corrected the inflation of p-values in standard association tests and showed that only a small subset of associations is directly linked to the disease. Direct associations had a much higher accuracy in separating cases from controls and pointed to immunomodulation, butyrate production, and the brain-gut axis as important factors in the inflammatory bowel disease.

## Introduction

Microbes are essential to any ecosystem be it the ocean or the human gut. The sheer impact of microbial processes has however been underappreciated until the advent of culture-independent methods to assess entire communities *in situ*. Metagenomics and 16S rRNA sequencing identified significant differences in microbiota among hosts, and experimental manipulations established that microbes could dramatically alter host phenotype [1–8]. Indeed, anxiety, obesity, colitis, and other phenotypes can be transmitted between hosts simply by transplanting their intestinal flora [9–13].

New tools and greater awareness of microbiota triggered a wave of association studies between microbiomes and host phenotypes. Microbiome wide association studies (MWAS) have been carried out for diabetes, arthritis, cancer, autism and many other disorders [14–23]. MWAS clearly established that each disease is associated with a distinct state of intestinal dysbiosis, but they often produced conflicting results and identified a very large number of associations both within and across studies [14, 19, 21, 23–26]. For example, a recent study on inflammatory bowel disease (IBD) reported close to 100 taxa associated with IBD [25], a number that is fairly typical [14]. Such long lists of associations defy simple interpretation and complicate mechanistic follow-up studies because one needs to examine the role of almost every species in the microbiota. In fact, one can argue that MWAS are most useful when they can identify a small network of taxa driving the disease.

Although extensive dysbiosis might reflect the multifactorial nature of the disease, it is also possible that MWAS detect spurious associations because their statistical methods fail to account for some important aspects of microbiome dynamics. One such aspect is the pervasive nature of microbial interactions: species compete for similar resources, rely on cross-feeding for survival, and even produce their own antibiotics [27–37]. Hence, microbial abundances must be correlated with each other, and even a simple change in host phenotype could manifest as collective responses by the microbiota. Traditional MWAS, however, completely neglect this possibility because they treat each change in abundance as an independent manifestation of altered host phenotype. As a result, MWAS cannot distinguish taxa directly linked to disease from taxa that are affected only through their interactions with other species.

The main conclusion of this paper is that realistic microbial interactions produce a large number of spurious associations. Many of these indirect associations can be removed by a simple procedure based on maximum entropy models from statistical physics, which can separate host effects from the microbial interactions. We dubbed this approach Direct Association Analysis, or DAA for short.

When applied to the largest MWAS on IBD, DAA shows that many of the previously reported associations could be explained by interspecific interactions rather than the disease. At the genus and species level, the direct associations include only *Roseburia*, *Faecalibacterium prausnitzii*, *Bifidobacterium adolescentis*, *Blautia producta*, *Turicibacter*, *Oscillospira*, *Eubacterium dolichum*, *Aggregatibacter segnis*, and *Sutterella*. Some of these associations are well-known, while others have received little attention in IBD research. The phenotypes of the taxa directly to disease suggest that immunomodulation, butyrate production, and the brain-gut interactions play important role in the etiology of IBD.

Compared to traditional MWAS, DAA corrected the inflation of p-values responsible for the large number of spurious associations and identified taxa most informative of the diagnosis. We found that directly associated taxa are much better at discriminating between cases from controls than an equally-sized subset of indirect associations. In fact, direct associations have the same potential to discriminate between health and disease as the entire set of almost a hundred associations detected by a conventional method.

## Results

Traditional MWAS detect species with significantly different abundances between case and control groups. Some changes in the abundances are directly associated with the disease while others are due to microbial interactions. The emergence of indirect changes in abundance is illustrated in Fig. 1A for a hypothetical network of five species. Only two species A and D are directly linked to the disease. However, strong interactions make the abundances of all five species differ between control and disease groups. For example, the mutualistic interaction between A and B helps B grow to a higher density following the increase in the abundance of A. The expansion of B in turn inhibits the growth of C and reduces its abundance in disease. Strong mutualistic, competitive, commensal, and parasitic interactions have been demonstrated in microbiota [27–37], and Fig. 1B shows that almost every species present in the human gut participates in a strong interaction. Thus, the propagation of abundance changes from directly-linked to other species could pose a significant challenge for MWAS. To test this hypothesis, we turned to a minimal mathematical model of microbiota composition.

**Figure 1.**
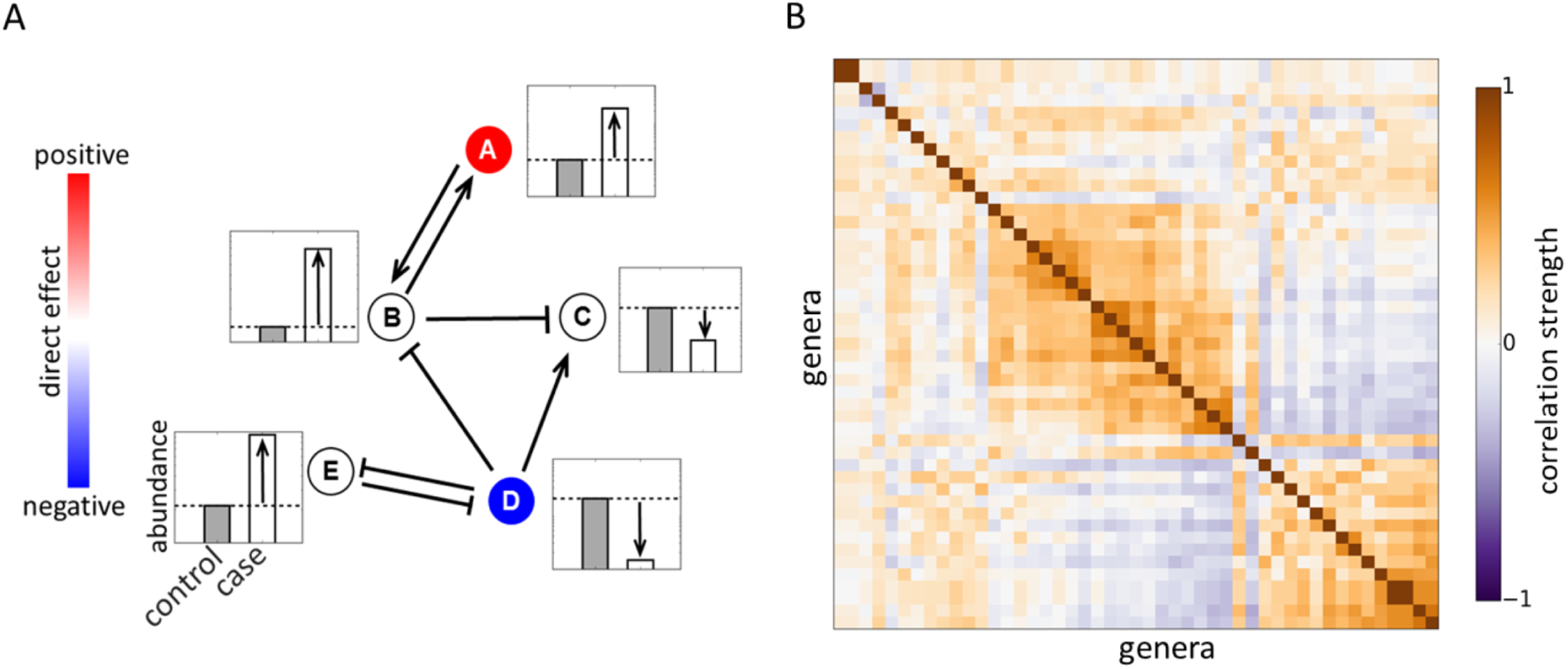
Microbial interactions generate spurious associations. **(A)** A hypothetical interaction network of five species together with their dynamics in disease. Only two species (shown in color) are directly linked to host phenotype. These directly-linked species inhibit or promote the growth of the other members of the community (shown with arrows). As a result, all five species have different abundances between case and control groups. **(B)** Microbial interactions are visualized via a hierarchically-clustered correlation matrix computed from the data in Ref. [21]. We used Pearson’s correlation coefficient between log-transformed abundances to quantify the strength of co-occurrence for each genus pair. Dark regions reflect strong interspecific interactions that could potentially generate spurious associations. See Tab. S1 for the list of 47 most prevalent genera included in the plot.

### Maximum entropy model of microbiota composition

A quantitative description of interspecific interactions and their effect on MWAS requires a statistical model of host-associated microbial communities. Ideally, such a model would describe the probability to observe any microbial composition, but the amount of data even in large studies is only sufficient to determine the means and covariances of microbial abundances. This situation is common in the analysis of biological data and has been successfully managed with the use of maximum entropy distributions [38]. These distributions are chosen to be as random as possible under the constraints imposed by the first and second moments. Maximum entropy models introduce the least amount of bias and reflect the tendency of natural systems to maximize their entropy [39]. In other contexts, these models have successfully described the dynamics of neurons, forests, flocks, and even predicted protein structure and function [40–44]. In the context of microbiomes, a recent work derived a maximum entropy distribution for microbial abundances using the principle of maximum diversity [45].

We show in the Supplemental information (SI) that the maximum entropy distribution of microbial abundances *P* ({*l*_*i*_}) takes the following form

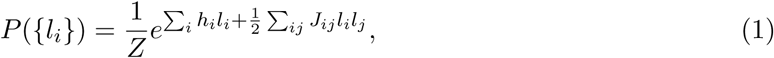

where *l*_*i*_ is the log-transformed abundance of species *i*, *h*_*i*_ represents the direct effect of the host phenotype on species *i*, and *J*_*ij*_ describes the interaction between species *i* and *j*; the factor of 1*/Z* is the normalization constant. The log-transformation of relative abundances alleviates two common difficulties with the analysis of the microbiome data. The first difficulty is the large subject-to-subject variation, which is much better captured by a log-normal rather than a Gaussian distribution; see Fig. S1, SI, and Ref. [25]. The second difficulty arises from the fact that the relative abundances must add up to one. This constraint is commonly known as the compositional bias because it leads to artifacts in the statistical analysis. The log-transformation is an essential step in most methods that account for the compositional bias [46–48], and, in the SI, we show that all of our conclusions are robust to the variation in the strength of the compositional bias.

### Testing for spurious associations in synthetic data

We obtained realistic model parameters from one of the largest case-control studies previously reported in Ref. [21]. The samples were obtained from mucosal biopsies of 275 newly diagnosed, treatment-naive children with Crohn’s disease (a subtype of IBD) and 189 matched controls. Microbiota composition was determined by 16S rRNA sequencing with about 30,000 reads per sample. From this data, we inferred the interaction matrix *J* and the typical changes in microbial abundances associated with the disease for 47 most prevalent genera (Methods and SI). Even though the number of data points significantly exceeds the number of free parameters in the model, overfitting could still be a potential concern. Overfitting, however, is unlikely to affect our main conclusions because they depend only on the overall statistical properties of *J* rather than on the precise knowledge of every interaction. In fact, none of our results changed when we analyzed only about half of the data set (Fig. 2). To improve the quality and robustness of the inference procedure, we also used the spectral decomposition of *J* to remove any interaction patterns that were not strongly supported by the data; see Methods and SI for further details.

To determine the effect of microbial interactions on conventional MWAS analysis, we generated synthetic data with a known number of direct associations. The data for the control group was used without modification from Ref. [21]. The disease group was generated using Eq. (5) with the same values of *h* and *J* as in the control group, except we modified the values of *h* for 6 representative genera (Tab. S2). We also generated two other synthetic data sets with smaller and larger effect sizes (Tab. S2). The results for all three data sets were very similar (SI).

The synthetic data was further subsampled to several sample sizes in order to simulate variation in statistical power between different studies. For an ideal method, the number of detected associations should increase with the cohort size, but eventually saturate once all 6 directly associated genera are discovered. In contrast to this expectation, the number of associations detected by the conventional approach increased rapidly with the sample size until almost all genera were found to be statistically associated with the disease in our synthetic data. At this point, traditional MWAS completely lost the power to identify the link between the phenotype and microbiota. Unbounded growth in the number of detections was also observed for the real data (Fig. 2C) suggesting that many previously reported associations between microbiota and IBD could be indirect.

Are spurious associations simply an artifact of our ability to detect even minute differences between cases and controls? Fig. 2B and 2D show that this was not the case. The median effect size declined only moderately with the number of associations, and most associations corresponded to about a factor of two difference in the taxon abundance. Thus, spurious associations are not weak and could not be discarded based on their effect size.

### Direct association analysis (DAA)

Fortunately, the maximum entropy model provides a straightforward way to separate direct from indirect associations. Since direct effects are encoded in *h*, MWAS should be performed on *h* rather than on *l*. This simple change in the statistical analysis correctly recovered 4 out of 6 directly associated taxa in the synthetic data and yielded no indirect associations even for large cohorts (Fig. 2A and S5). Similarly good performance was found for the two other synthetic data sets (Fig. S7). For the IBD data, DAA also identified a much smaller number of associations compared to traditional MWAS analysis and showed clear saturation at large sample sizes (Fig. 2B). Direct associations with IBD are summarized in Fig. 3 at the genus and species levels, and the entire phylogenetic tree of direct associations is shown in Fig. S2 and Tabs. S3 and S4.

To demonstrate that DAA isolates direct effects from collective changes in the microbiota, we examined the p-value distribution in this method. The distribution of p-values is commonly used as a diagnostic tool to test whether a statistical method is appropriate for the data. In the absence of any associations, p-values must follow a uniform distribution because the null hypothesis is true [54]. A few strong deviations from the uniform distribution signal true associations [55]. In contrast, large departures from the uniform distribution typically indicate that the statistical method does not account for some properties of the data, for example, population stratification in the context of genome wide association studies [56, 57]. Figure 4A compares the distribution of p-values for DAA and a conventional method in MWAS. Consistent with our hypothesis that interspecific interactions cannot be neglected, conventional analysis generates an excess of low p-values and, as a result, a large number of potentially indirect associations. In contrast, the distribution of p-values from DAA matches the expected uniform distribution and, thus, provides strong support for our method.

Finally, we show that indirect association excluded by DAA do not reduce the predictive power of microbiome data. Supervised machine learning such as random forest [58, 59], support vector machine [60], and sparse logistic regression [61–63] were used to classify samples as cases or controls based on their microbiota profile. We found good and identical performance of the classifiers trained either on all taxa detected by conventional MWAS or on a much smaller subset of direct associations detected by DAA (Fig. 4B). Moreover, the DAA-based classifier showed significantly better performance compared to a classifier trained on an equal number of randomly-selected indirect associations (Fig. 4B). Thus, DAA reduces the number of associations without losing any information on the disease status and selects taxa with the greatest potential to distinguish health from disease.

**Figure 2.**
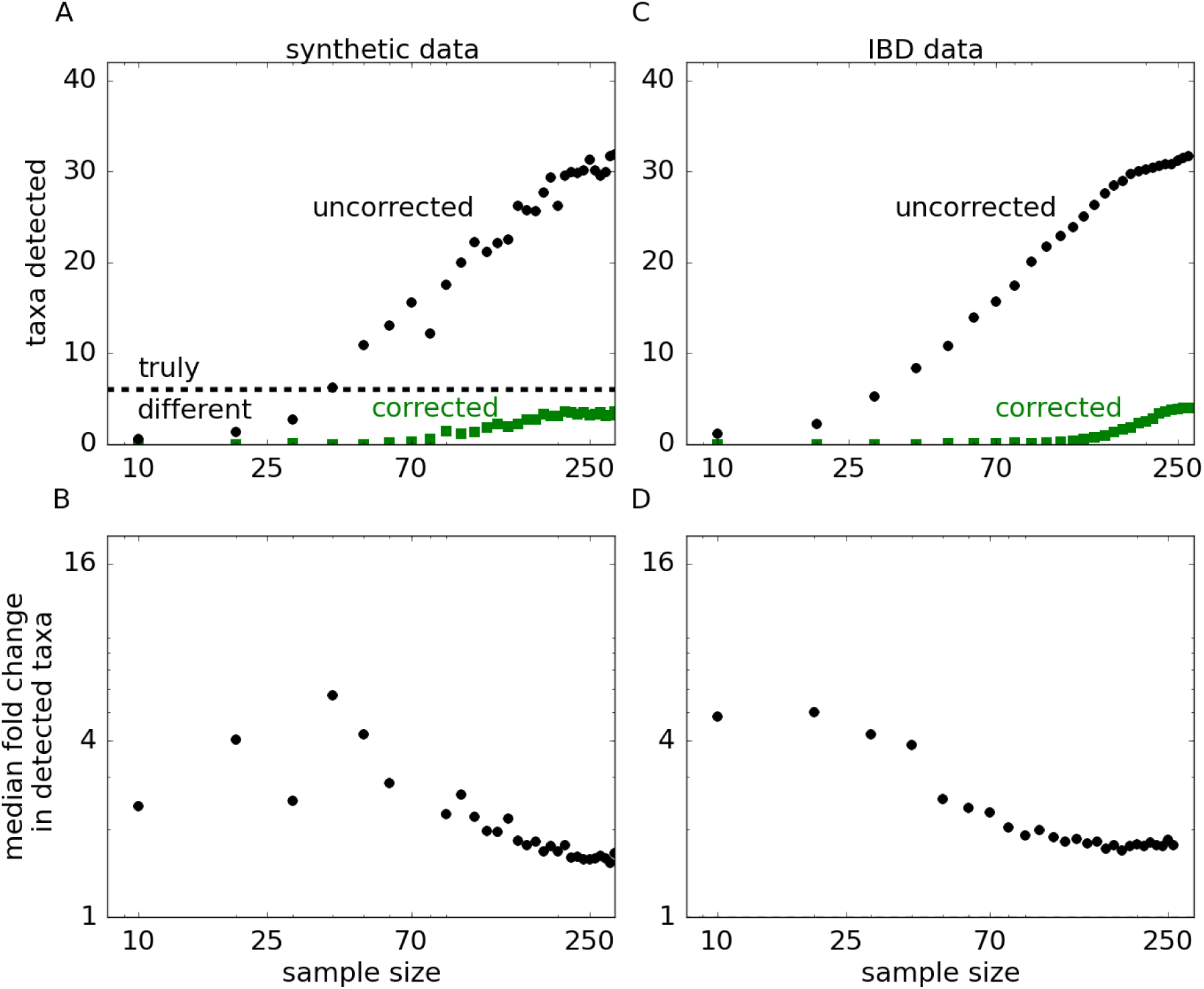
Signatures of indirect associations in synthetic and IBD data sets. The synthetic data set was generated to match the statistical properties of the IBD data set from Ref. [21], but with a predefined number of 6 directly associated taxa. **(A)** In synthetic data, DAA identifies no spurious association and detects 4 out of 6 directly associated genera. All 6 genera and no false positives are detected when the sample size is increased further (Fig. S5). In sharp contrast, a large number of spurious associations is observed for metrics that rely on changes in abundance between cases and controls and do not correct for microbial interactions. The number of false positives grows rapidly with statistical power until all taxa are reported as significantly associated with the disease (Fig. S5). **(B)** All spurious associations show substantial differences between cases and controls and, therefore, cannot be discarded based on their effect sizes. To quantify the effect size, we estimated the magnitude of the fold change for each genus. Specifically, we first computed the difference in the mean log abundance between cases and controls and then exponentiated the absolute value of this difference. The plot shows how the median effect size for significantly associated genera depends on the sample size. Larger samples sizes result in much higher number of associations, but only a small drop in the typical effect size. (**C**) and (**D**) are the same as (A) and (B), but for the IBD data set. The results are consistent between the two data sets suggesting that most associations detected by traditional MWAS are spurious. The complete list of indirect associations inferred from the IBD data set is shown in Tab. S5 and the results for different synthetic data sets are shown in Fig. S7.

**Figure 3.**
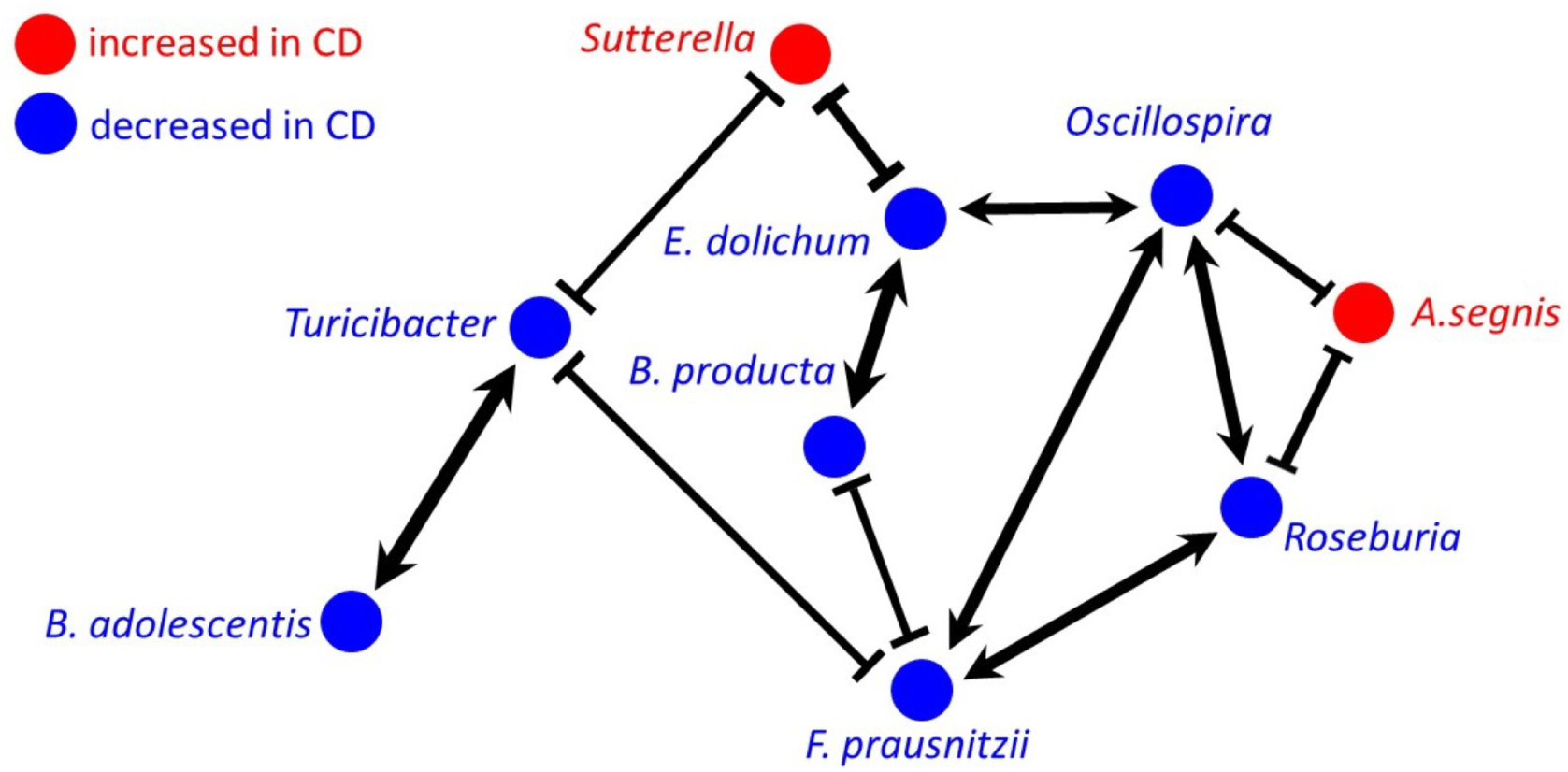
Network of direct associations with Crohn’s Disease. Five species and four genera were found to be significantly associated with Crohn’s Disease (*q <* 0.05) after correcting for microbial interactions. The links correspond to significant interactions (*q <* 0.05) between the taxa with *J*_*ij*_ *>* 0.27 or *J*_*ij*_ *<* -0.15; the width of the arrows reflects the strength of the interactions. Note that DAA controls for the fact that two species could be correlated because both interact with a third species, but not with each other (Fig. 1A). Thus, the network shows only direct interactions between the taxa. Compared to the correlation matrix in Fig. 1B, the interaction network has both mutualistic and inhibitory links, which suggests that the microbial community might have several stable states corresponding to distinct modes of dysbiosis [30, 49–53]. For comparison, the correlation-based network for directly associated taxa is shown in Fig. S3. A complete summary of correlations and interactions for all species pairs is provided in Tab. S6.

## Discussion

The primary goal of MWAS is to guide the study of disease etiology by detecting microbes that have a direct effect on the host. These direct effects could be very diverse and include secretion of toxins, production of nutrients, stimulation of the immune system, and changes in mucus and bile [64, 65]. In addition to the host-microbe interactions, the composition of microbiota is also influenced by the interspecific interactions among the microbes such as competition for resources, cross-feeding, and production of antibiotics. In the context of MWAS, microbial interactions contribute to indirect changes in microbial abundances, which are less informative of the disease mechanism and are less likely to be valuable for follow-up studies or in interventions. Here, we estimated the relative contribution of indirect associations to MWAS and showed how to isolate direct from indirect assoctiations.

Our main result is that interspecific interactions are sufficiently strong to generate detectable changes in the abundance of many microbes that are not directly linked to host phenotype. As a result, conventional approaches to MWAS detect a large number of spurious associations and produce inflated p-values that do not match their expected distribution (Fig. 4A). These challenges are resolved by Direct Association Analysis (DAA), which uses maximum entropy models to explicitly account for interspecific interactions. We applied DAA to a large data set of pediatric Crohn’s disease and found that it restores the distribution of p-values and substantially simplifies the pattern of dysbiosis while retaining full classification power of a conventional MWAS.

**Figure 4.**
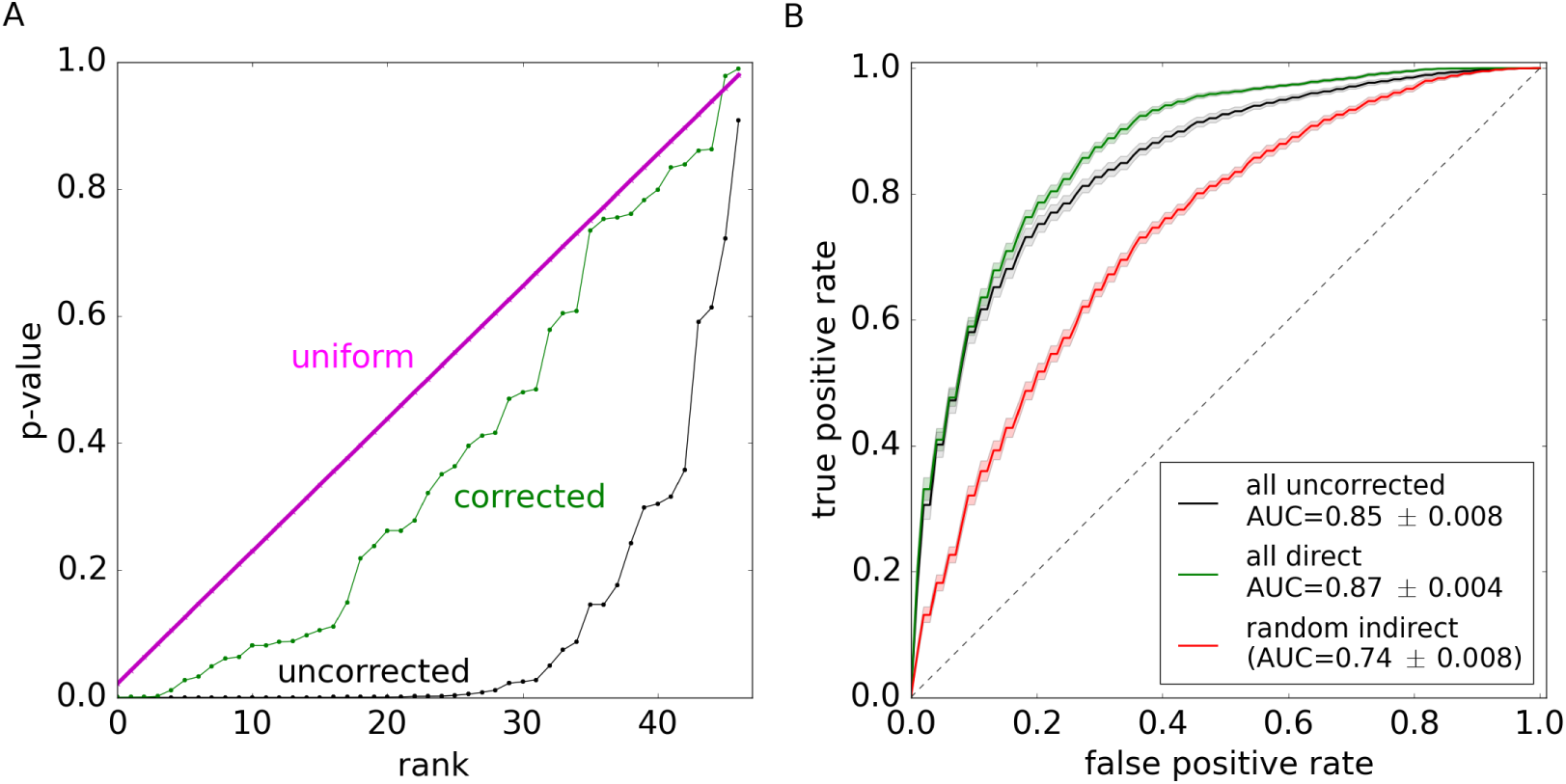
Direct associations analysis corrects p-value inflation and retains diagnostic accuracy. **A)** The distribution of p-values in DAA closely follows the expected uniform distribution. Without correcting for microbial interactions, the same analysis yields an excess of low p-values, a signature of indirect associations. For both methods, p-values were computed using a permutation test. The expected uniform distribution was obtained by sampling from a generator of random numbers. The ranked plots of p-values visualize their cumulative distribution functions; this is a variant of a Q-Q plot. **(B)** Direct associations are a small subset of all associations with IBD, yet they retain full power in classifying samples as cases or controls. In contrast, the classification power is substantially reduced for an equally-sized subset of randomly-chosen indirect associations. In each case, we used sparse logistic regression to train a classifier on 80% of the data and tested its performance on the remaining 20% (Methods). The shaded regions show one standard deviation obtained by repeated partitioning the data into the training and validation sets. Identical results were obtained with a random forest [58, 59] and support vector machine [60] classifiers (Fig. S4).

The relatively simple dysbiosis identified by DAA in IBD has strong support in the literature and offers interesting insights into disease etiology. Four of the taxa identified by our method have a well-established role in IBD: *B. adolescentis*, *F. prausnitzii*, *B. producta*, and *Roseburia*. They have been repeatedly found to have lower abundance in both Crohn’s disease and ulcerative colitis [66–73], and several studies have demonstrated their ability to suppress inflammation and alleviate colitis [69, 74–78]. *Bifidobacterium* species occupy a low trophic level in the gut and ferment complex polysaccharides such as fiber [79, 80]. Fermentation products include lactic acid, which promotes barrier function, and maintains a healthy, slightly acidic environment in the colon [81]. Due to these properties *Bifidobacterium* species are commonly used as probiotics [79]. *F. prausnitzii*, *Blautia producta* and *Roseburia* occupy a higher trophic level and ferment the byproducts of polysaccharide digestion into short-chain fatty acids (SCFA), which are an important energy source for the host [68,69, 82, 83].

The ability of DAA to detect taxa strongly associated with IBD is reassuring, but not surprising. What is surprising is that many strong associations are classified as indirect by our method. For example, *Roseburia* and *Blautia* are the only genera of *Lachnospiraceae* that DAA finds to be directly linked to the disease. In sharp contrast, traditional MWAS report seven genera in this family that are strongly associated with IBD [25]. All seven genera are involved in SCFA metabolism, but their specializations differ. Species in *Blautia* genus are major producers of acetate, a SCFA that is commonly involved in microbial crossfeeding [84, 85]. In particular, many species extract energy from acetate by converting it into butyrate, another SCFA that plays a major role in gut health by nourishing colonocytes and regulating the immune function [82, 85]. *Roseburia* genus specializes almost exclusively in the production of butyrate and acts as a major source of butyrate for the host [82, 86]. Thus, our findings suggest that butyrate production plays an important role in IBD etiology and that the dysregulation of this process is directly linked to the depletion of *Roseburia* and possibly *Blautia*.

The important role of butyrate is further supported by our detection of *E. dolichum* and *Oscillospira*, which are known to produce butyrate [87–89]. The latter taxon has not been detected in three independent analyses of this IBD data set [21, 25, 90] presumably because its involvement was masked by indirect associations and interactions with other microbes. Indeed, several other studies found that *Oscillospira* is suppressed in IBD [91],[92]. *Oscillospira* was also found to be positively associated with leanness and negatively associated with the inflammatory liver disease [93–95]. The interactions between *Oscillospira* and the host appears to be quite complex and involve the consumption of host-derived glycoproteins including mucin, production of SCFA, and modulation of bile-acid metabolism [89, 96, 97]. The latter interaction was suggested to be a major factor in the protective role of *Oscillospira* against infections with *Clostridium difficile* [96, 98, 99].

The final taxon that was suppressed in IBD is *Turicibacter*. This genus is not very well characterized, and few MWAS studies point to its involvement in IBD [21, 25, 100]. Two studies in animal models, however, directly looked into the connection between IBD and *Turicibacter* [101, 102]. The first study found that iron limitation eliminates colitis in mice while at the same time restoring the abundance of *Turicibacter*, *Bifidobacterium*, and four other genera [101]. The second study study identified *Turicibacter* as the only genus that is fully correlated with immunological differences between mice resistant and susceptible to colitis: high abundance of *Turicibacter* in the colon predicted high levels of MZ B and iNK T cells, which are potent regulators of the immune response [102]. Moreover, *Turicibacter* was the only genus positively affected by the reduction in CD8^+^ T cells. Thus, our method identified a taxon that is potentially directly linked to IBD via the modulation of the immune system.

Perhaps the most unexpected finding was our detection of *Aggregatibacter* and *Sutterella* as the only genera increased in disease compared to 26 positive associations detected by the previous analysis [25]. All other associations were classified as indirect even though they often corresponded to much more significant changes in abundance between IBD and control groups. Thus, our results indicate that expansion of many taxa including opportunistic pathogens is driven by their interactions with the core IBD network shown in Fig. 3. One possibility is that the dysbiosis of the symbiotic microbiota makes it less competitive against other bacteria and opens up niches that can be colonized by opportunistic pathogens. The other, less explored possibility, is that commensal microbiota can not only protect from pathogens, but also facilitate their invasion, a phenomenon that has been recently demonstrated in bees [103].

Little is known about the specific roles that *Aggregatibacter* and *Sutterella* play in IBD, and more generally in gut health. *Aggregatibacter* is a common member of the oral microbiota that thrives in local infections such as periodontal disease and bacterial vaginosis [104–106]. The high abundance of *Aggregatibacter* is also associated with an increased risk of IBD recurrence [107]. *Sutterella*, on the other hand lacks overt pathogenicity, and MWAS produced inconsistent findings [108–114] on its involvement in IBD. Some studies reported that *Sutterella* is increased in patients with good outcomes [21, 111] while other studies found positive or no association between *Sutterella* and IBD [25, 109, 112–114]. Experimental investigations showed that *Sutterella* lacks many pathogenic properties; in particular, it does not induce a strong immune-response and has only moderate ability to adhere to mucus [113, 114]. Further, *Sutterella* strains from IBD and control patients showed no phenotypic differences in metabolomic, proteomic, and immune response assays [114]. Nevertheless, *Sutterella* is strongly associated with worse behavioral scores in children with autism spectrum disorder and Down syndrome [19, 20, 115]. Therefore, the direct link between *Sutterella* and IBD could involve the gut-brain axis.

In summary, we found a small number of taxa can explain extensive dysbiosis in IBD and accurately predict disease status. Directly associated taxa include strains with dramatically different abilities to trigger colitis and are specifically targeted by the immune system of patients and animals with IBD [12]. Previous studies of these taxa point to facilitated colonization by pathogens, butyrate production, immunomodulation, bile metabolism, and the gut-brain axis as the primary factors in the etiology of IBD.

Many disorders are accompanied by substantial changes in host microbiota, but our work shows that only a small subset of these changes could be directly related to the disease. Similarly, only a handful of taxa could drive the dynamics of ecosystem-level changes in the environment. To untangle the complexity of such dysbioses, it is important to account for microbial interactions using mechanistic or statistical methods. Direct association analysis is a simple statistical approach based on the principle of maximum entropy. It can be applied to any microbiome data set that is sufficiently large to infer interspecific interactions.

## Methods

The data used in this study was obtained from Ref. [21], which reported changes in the microbiome of newly-diagnosed, treatment-naive children with IBD compared to controls. This data was recently analyzed in Ref. [25], and we followed all the statistical procedures adopted in that study to enable direct comparison of the results. Specifically, we used a permutation test on mean log-transformed abundances to determine the statistical significance of an association.

All computation was carried out in Python environment. We used `scikit-learn 0.15.2` [116] for hierarchical clustering and to build the supervised classifiers used in Fig. 4B of the main text and Fig. S3. The variance in the accuracy of classification was evaluated through 5-fold stratified cross-validation with 100 random partitions of the data into the training and validation sets. For all findings, statistical significance was evaluated with Fisher’s exact test (permutation test) with 10^6^ permutations. False discovery rate was controlled to be below 5% following Benjamini-Hochberg method [54].

To fit the maximum entropy model to the data, we first computed the mean log-abundance for each genus *m*_*i*_ and the covariance in the log-transformed abundances *C*_*ij*_. The interaction matrix was computed as *J* = *C*^*-1*^ by performing singular value decomposition [117] and removing all singular values that were comparable to the amount of noise present in the data. The host effects were computed as *h* = *J m*. See Supplementary Methods for further details.

## Acknowledgements

This work was supported by a grant from the Simons Foundation (#409704, Kirill S. Korolev) and by the startup fund from Boston University to Kirill S. Korolev. Vivek Ramanan was also supported by NSF grant DBI-1559829 through BRITE Bioinformatics REU Program at Boston University. Rajita Menon was partly supported by the Graduate Fellowship from the Rafik B. Hariri Institute for Computing and Computational Sciences & Engineering. Simulations were carried out on Shared Computing Cluster at Boston University.

## Supplementary information for “Interactions between species introduce spurious associations in microbiome studies”

### Model of community composition

Here we describe a mathematical model of community composition, that we use to correct for microbial interactions in microbiome-wide association studies.

#### Log-transformation of abundances

The environment within a host is constantly changing due to variations in diet, immune response, phage activity and other factors. As a result, microbial growth rates should be highly variable and produce multiplicative fluctuations in the community composition, which are better captured on logarithmic rather than on linear scale. Indeed, the abundances of many gut species follow a log-normal distribution (Fig. S1), and recent work shows that a log-transformation of abundances increases the power and quality of microbiome studies [25]. Therefore, we chose to carry out all of the analysis and modeling on natural logarithms of relative abundances computed with a pseudocount of one read. For simplicity, we refer to these quantities as abundances in the following and denote them as *l*_*i*_ with the subscript identifying the species under consideration.

#### Maximum entropy models

Microbiota composition is highly variable among people in both health and disease [25] and needs to be described via a multivariate probability distribution *P* ({*l*_*i*_}). The amount of data in a large microbiome-wide association study, however, is sufficient to reliably determine only the first and second moments of *P* ({*l*_*i*_}). This situation is common in the analysis of biological data and has been successfully managed with the use of maximum entropy distributions [38]. These distributions are chosen to be as random as possible under the constraints imposed by the first and second moments. Maximum entropy models introduce the least amount of bias and reflect the tendency of natural systems to maximize their entropy. In other contexts, these models have successfully described the dynamics of neurons [40], forests [41], and flocks [42], and even predicted protein structure [43] and function [44]. In the context of microbiomes, a recent work derived a maximum entropy distribution for microbial abundances using the principle of maximum diversity [45].

Let us denote abundance means and covariances computed from the data by the vector *m* and matrix *C* respectively. The constraints on the maximum entropy distribution are then expressed as

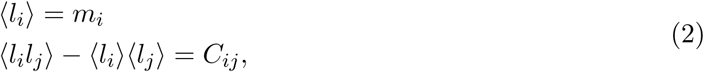

and the maximum entropy distribution takes the following form

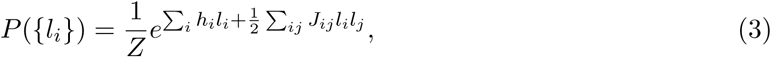

which is known as the Ising model in statistical physics. The variables *h*_*i*_ and *J*_*ij*_ arise as Lagrange multipliers for the first and second moment constraints during entropy maximization. In statistical physics, they describe local magnetic fields that align spins *l*_*i*_ and interactions between spins *l*_*i*_ and *l*_*j*_. The constant *Z*, known as the partition function, ensures that the distribution is normalized:

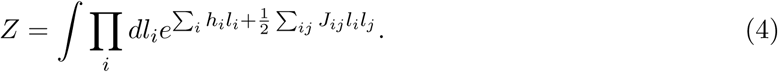

#### Host effects vs. species interactions

To interpret this maximum entropy distribution in terms of biologically relevant factors such as microbial interactions and properties of the host, we can rewrite equation (5) as follows

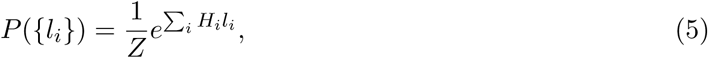

where

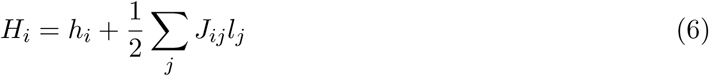

describe the quality of the local environment for species *i*: the higher *H*_*i*_, the more abundant the species. The quality of the environment can be decomposed into external variables such as temperature or metabolite concentrations *V*_*α*_ and the species’ response to these variables *R*_*iα*_ as

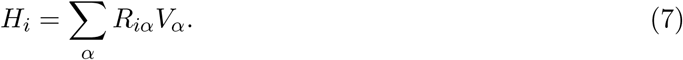

We can further decompose the external variables *V*_*α*_ into host factors 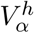 and influences of other species, e.g., due to metabolite secretion or production of antibiotics:

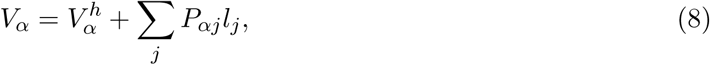

where *P*_*αj*_ describes the influence of microbe *i* on variable *α*.

Upon combining equations (7) and (8), we can express *H*_*i*_ as

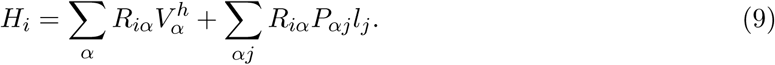

Comparison of this equation to equation (6) shows that we can identify 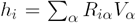 with the direct effects of the host and 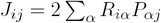 with the interactions among the microbes.

### Inference of model parameters

Here we describe the procedure of learning the parameters of the maximum entropy model from the data. Our approach closely follows that of Refs. [38], [43] and [44].

#### Relating h and J to m and C

To infer model parameters *h*_*i*_ and *J*_*ij*_, we need to relate them to empirical observations such as the means and covariances of the abundances. These relationships can be conveniently obtained from the derivatives of the partition function, which is the standard approach in statistical physics. Indeed, the mean abundances can be expressed as

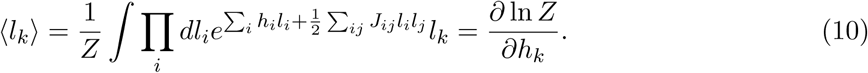

A similar relationship holds for the covariance matrix:

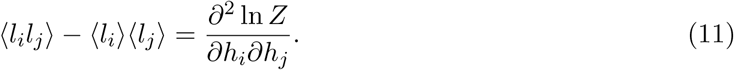

To complete the calculation, we need to compute the partition function defined by equation (4). The result reads

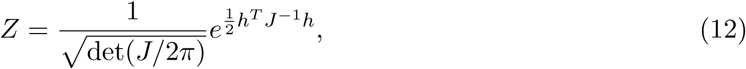

where symbols without indexes are treated as vectors or matrices.

From equation (12), we immediately find that

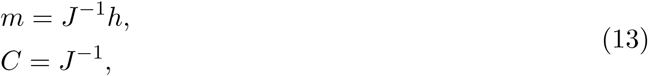

which can be inverted to obtain

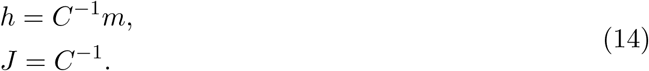

#### Inverting the covariance matrix

It is clear from equation (14) that the key step in obtaining the model parameters is the inversion of the covariance matrix. However, this matrix is likely to be degenerate or ill-conditioned because of the insufficient amount of data or very strong correlations between microbial abundances. To overcome this difficulty, we computed a pseudoinverse of *C* as described in the following sections. Briefly, we used singular value decomposition [117] of *C* in terms of two orthogonal matrices *U* and *V* and a diagonal matrix Λ:

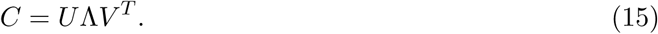

Some diagonal elements of Λ were small and comparable to the levels of noise (or uncertainty), so we set the corresponding elements of Λ^-1^ to zero. Specifically, 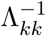 was set to zero for all *k* such that Λ*_kk_ < λ*_min_, where *λ*_*min*_ was a predetermined threshold. A regular inverse 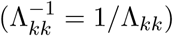 was used for the rest of the elements. The robustness of the results to the variation in the threshold *λ*_min_ is discussed in the section on data analysis. This procedure ensured that we do not infer large changes in host fields *h* due to fluctuations in the estimate of ⟨*l*⟩. The inverse of *C* was then computed as *C*^-1^ = *V* Λ^-1^*U^T^*, where we used the fact that the inverse of an orthogonal matrix is its transpose.

### Origin of spurious associations and Direct Associations Analysis

#### Microbial interactions introduce spurious associations

In microbiome-wide association studies, we are typically interested in the changes in microbial abundances Δ*m* between two groups of subjects. From equation (13), we can relate Δ*m* to the changes in the phenotype of the host Δ*h*:

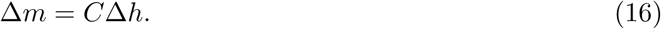

This formula clearly illustrates the origin of spurious associations. Imagine that there is a small number of species directly linked to host phenotype, i.e. Δ*h* is a sparse vector. Because *C* is a dense matrix (see Fig. 1b in the main text), equation (16) predicts that Δ*m* is dense, i.e. the abundances of most species are affected. The sizes of these effects are variable and depend on the magnitude of the off-diagonal elements of *C*. Except for the strongly interacting species, the largest changes in *m* are likely to mirror the largest changes in *h* and result in significant associations. In large samples, however, smaller effects become detectable that could either reflect small direct effects or the secondary, indirect effects due to microbial interactions. As a result, the number of associations grows with the sample size, and the relationship between associated species and host phenotype becomes obscured. Fig. 2 in the main text presents evidence for a large number of spurious associations in both synthetic and real data.

#### Removing indirect associations

Equation (16) offers as straightforward way to correct for microbial interactions and separate direct from indirect associations. Indeed, for each species, we can compute the corresponding change in the host field as

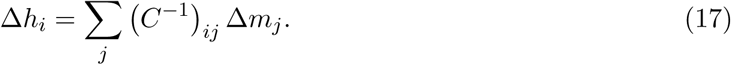

The statistical significance of this change can be determined via the permutation test followed by the Benjamini-Hochberg procedure to correct for multiple hypothesis testing [54].

### Generation of synthetic data

Here, we describe how we generated the synthetic data shown in Fig. 2A of the main text. This data was generated to evaluate the likelihood of spurious associations in MWAS. We introduced a known number of direct associations, but ensured that all other properties of the data correspond to that of the human gut microbiota.

The data for the control group were directly subsampled from the IBD data set. To generate the data for the disease group, we first inferred the covariance matrix using the entire data set and the mean abundances using just the control group. Then, equation (13) was used to compute *h*. These values of *h* described normal microbial abundances in subject without IBD. To introduce a difference between cases and controls, we modified the values of *h* for 6 randomly chosen species by 10% - 40%; these are typical changes in *h* identified by DAA. Finally, we computed the expected microbial abundance using equation (13) and then sampled from a multivariate normal distribution with these means and the covariance matrix defined above.

We also tested that our conclusions hold for other diseases with potentially different effect sizes. Specifically, we repeated the analysis in Fig. 2A for two other synthetic data sets: one with smaller and one with larger effect sizes. The results are qualitatively similar to what we reported in the main text and are shown in Fig. S7. The values of the effect sizes are given in Tab. S2.

### Data analysis

For correlation analysis, we used Pearson correlation coefficient for log-transformed abundances.

For logistic regression classifier, we used L1 penalty to ensure sparseness and generalizability. In all classifiers default parameters were used in scikit-learn version 0.17.2.

For hierarchical clustering of the correlation matrix, we used the Nearest Point Algorithm method of the linkage function in scipy with a correlation distance metric.

#### Threshold for matrix inversion

For our analysis of the IBD and synthetic data sets we set *λ*_min_ to 0.01. To test whether our results are robust to the value of the threshold, we varied the number of eigenvalues of Λ*^-1^* not set to zero; see Fig. S8. When only a few eigenvalues where included, DAA detected a large number of associations because many taxa were perfectly correlated, and it was impossible to distinguish direct from indirect associations. As the number of included eigenvalues increased, the performance of DAA improved and reached a plateau. In this plateau region, the results were largely insensitive to the value of the threshold used.

#### Compositional effects

Microbiota composition is usually quantified by relative abundances to eliminate the variation in the total number of sequencing reads. Although the total number of reads depends on the total abundance of microbes, variation in sample preparation and other factors also contribute and thereby make the inference of absolute microbial abundances nearly impossible. Because the relative abundance add up to one, microbiome data is compositional, which leads to spurious correlations and other statistical biases [46–48]. These biases are largely removed by the log-transformation and we find no evidence that they significantly affect our conclusions including the results of DAA. This can be seen from Fig. S6, which compares the analysis done on relative abundances to the analysis done on unnormalized counts. Both analyses identify about the same number of associations (and the same taxa) using either traditional MWAS or DAA. Note that a very strong compositional bias would make all taxa associated with the disease simply because the change in the relative abundance of one taxon necessarily changes the abundance of all other taxa. Such strong compositional bias is not observed in either IBD data set or in synthetic data with fewer than 5000 samples. Finally, we note that our synthetic data has the same amount of compositional bias as in the IBD data. For both data sets, the top 10 most abundant taxa account for 80% of the reads. Compositional effects could be stronger in less diverse habitats with lower species evenness compared to the gut.

### Computer code

We include here the link to computer code that loads the data and outputs all figures and tables: https://github.com/rajitam/DAA-figures-and-tables

**Figure S1.**
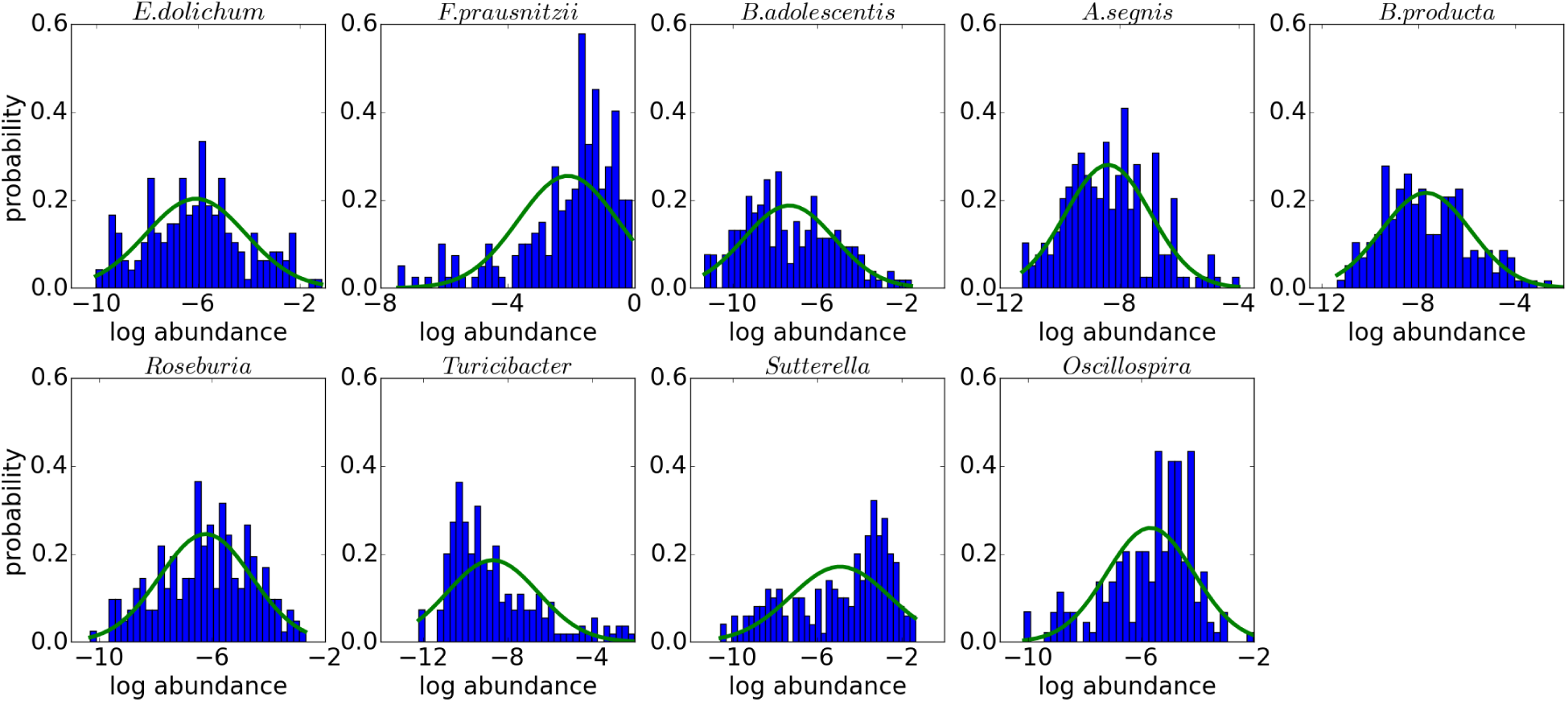
Microbial abundances follow the log-normal distribution. The histograms show probability distributions of the relative log-abundance for the species and genera detected by DAA. The best fit of a Gaussian distribution is shown in green.

**Figure S2.**
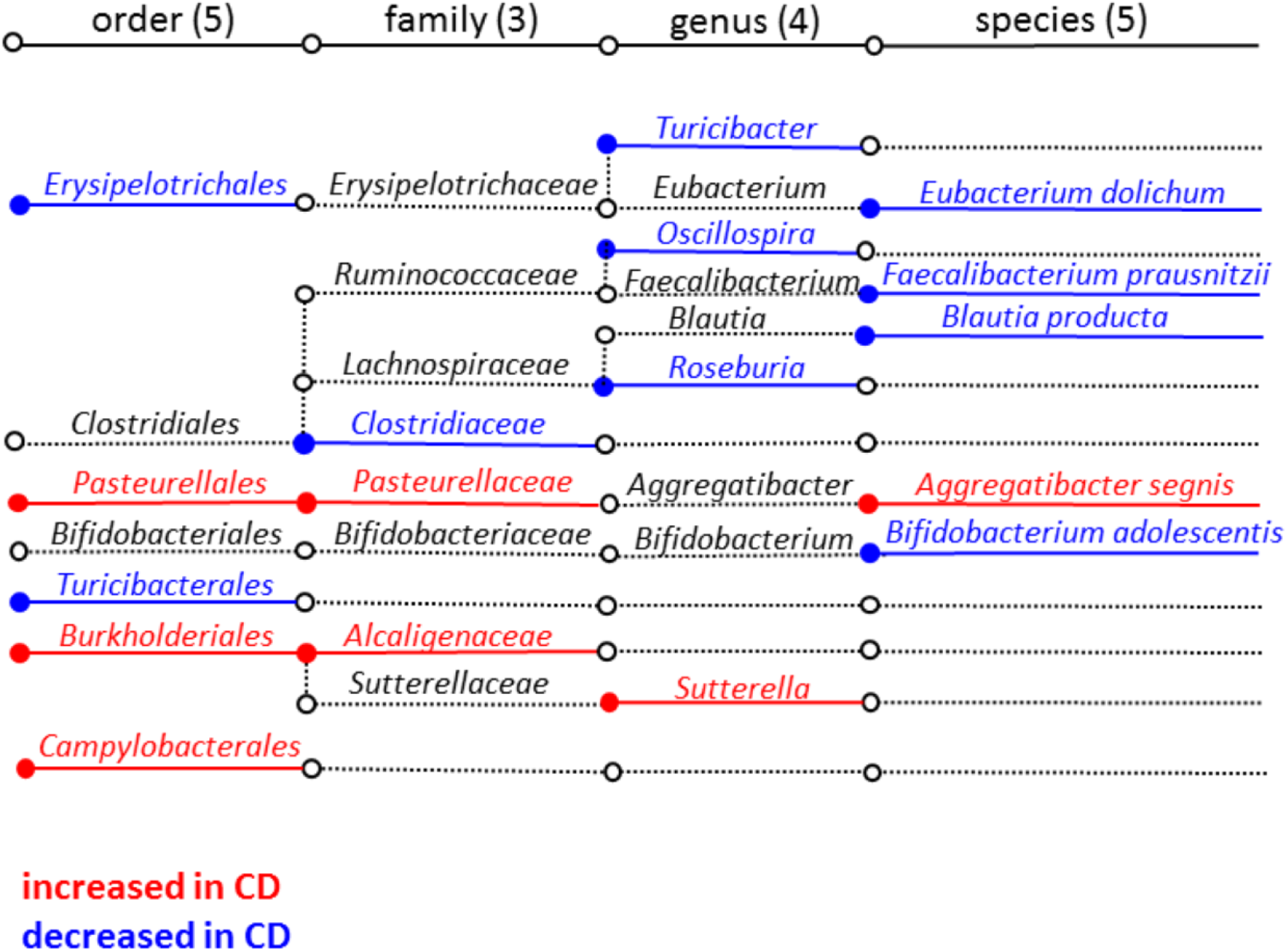
Taxa directly associated with Crohn’s disease. Note that the Green Genes database [118] used in QIIME [119] places Turicibacter under Erysipelotrichales and has a unique order of Turicibacterales. This apparent inconsistency may reflect insufficient understanding of Turicibacter phylogeny. The effect sizes and statistical significance are summarised in Tab. S3 and compared between DAA and conventional MWAS in Tab. S4.

**Figure S3.**
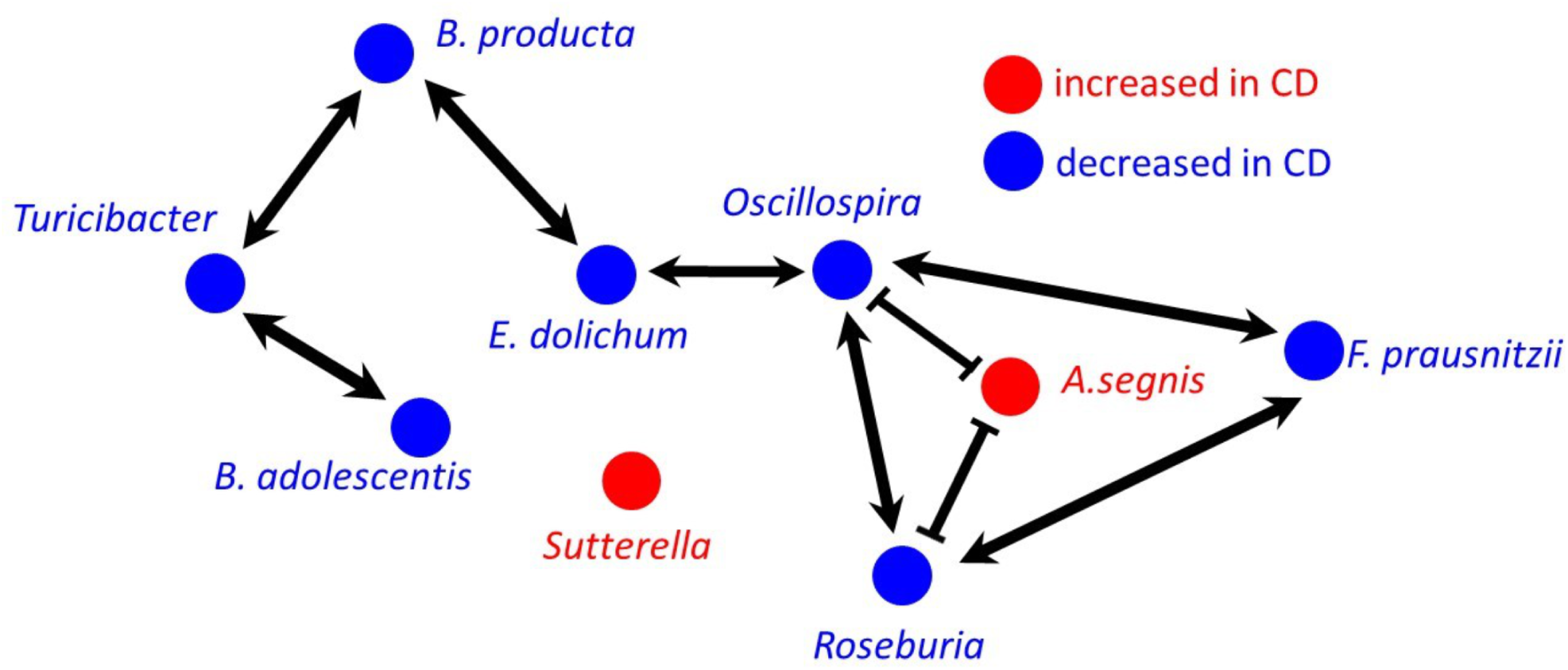
The network based on the correlation coefficient between log transformed abundances. We plotted the correlation based network for the species detected by DAA. Note the similarities and differences with the interaction network shown in Fig. 3 of the main text. Only the links with the correlation coefficient greater than 0.27 or lower than -0.15 are shown, and all links are statistically significant (*q <* 0.05). All correlation coefficients and direct interactions are summarized in Tab. S6 for the genera and species detected by DAA.

**Figure S4.**
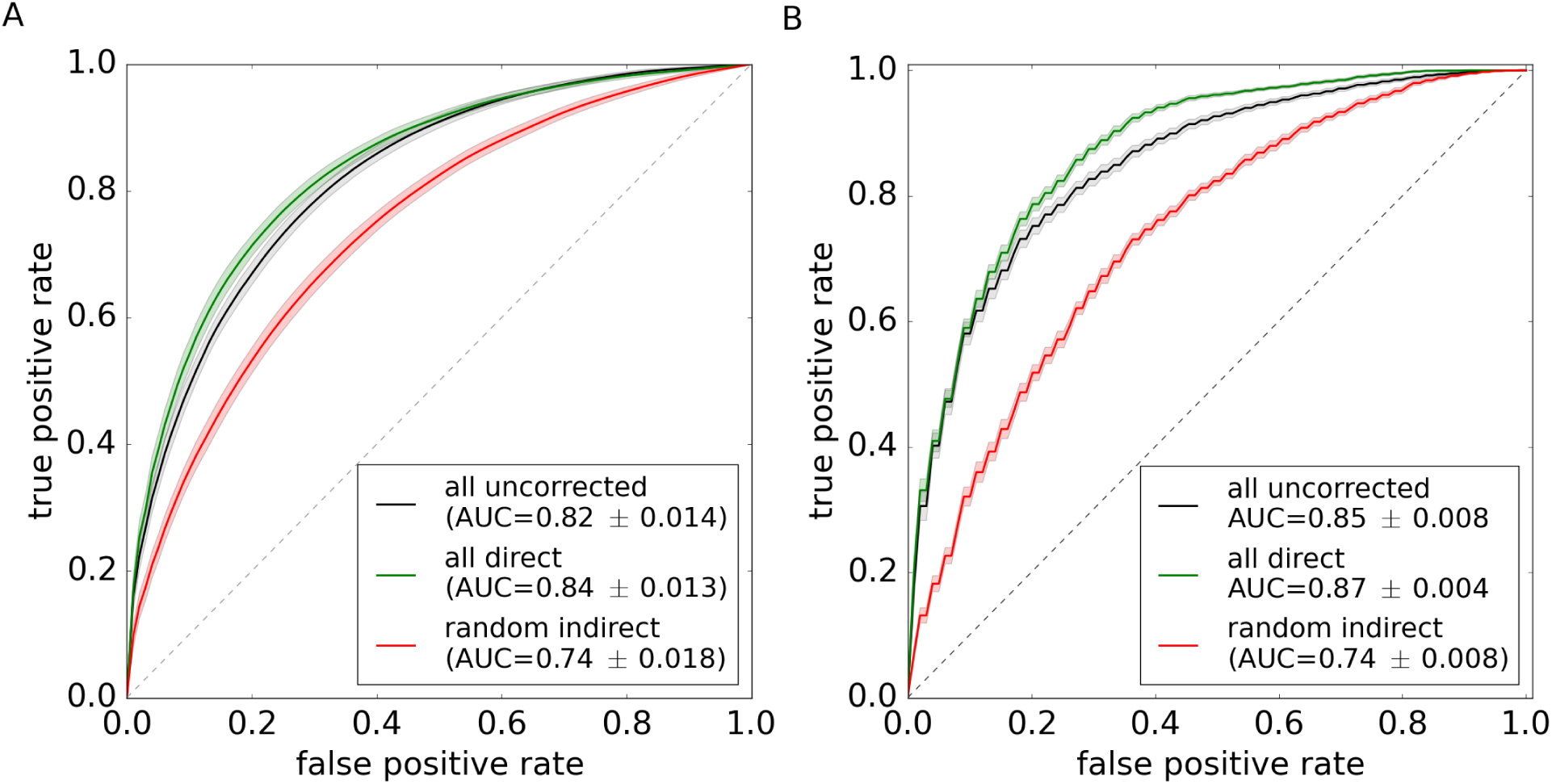
Direct associations retain full diagnostic power. The same as Fig. 4B of the main text, but for two other classifiers: random forest [58, 59] in **(A)** and support vector machine [60] in **(B)**.

**Figure S5.**
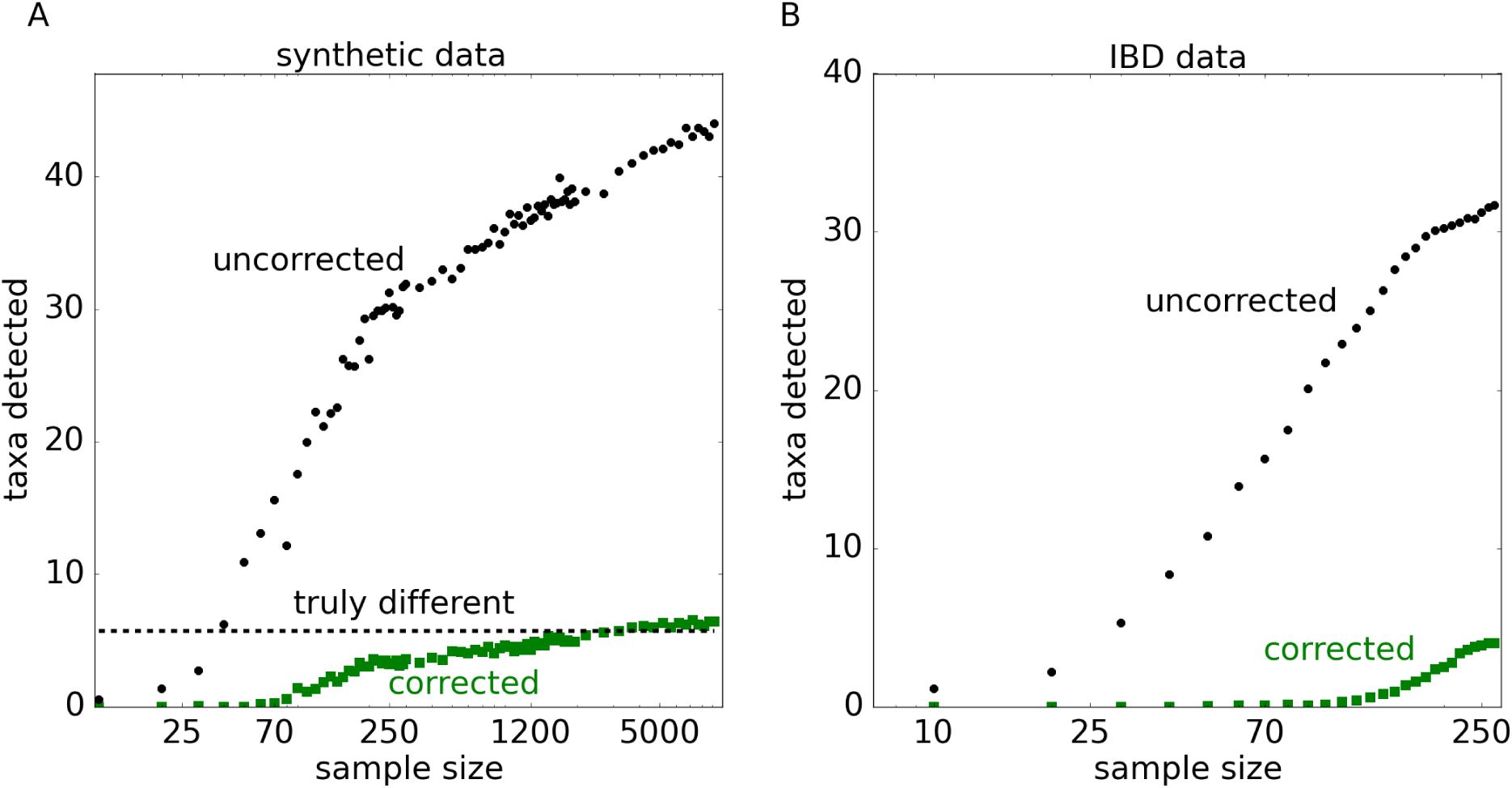
DAA detects all directly associated taxa in synthetic data with enough samples. The same as Fig. 2A, but with the x-axis extended to large sample sizes. Note that DAA recovers all 6 directly associated taxa.

**Figure S6.**
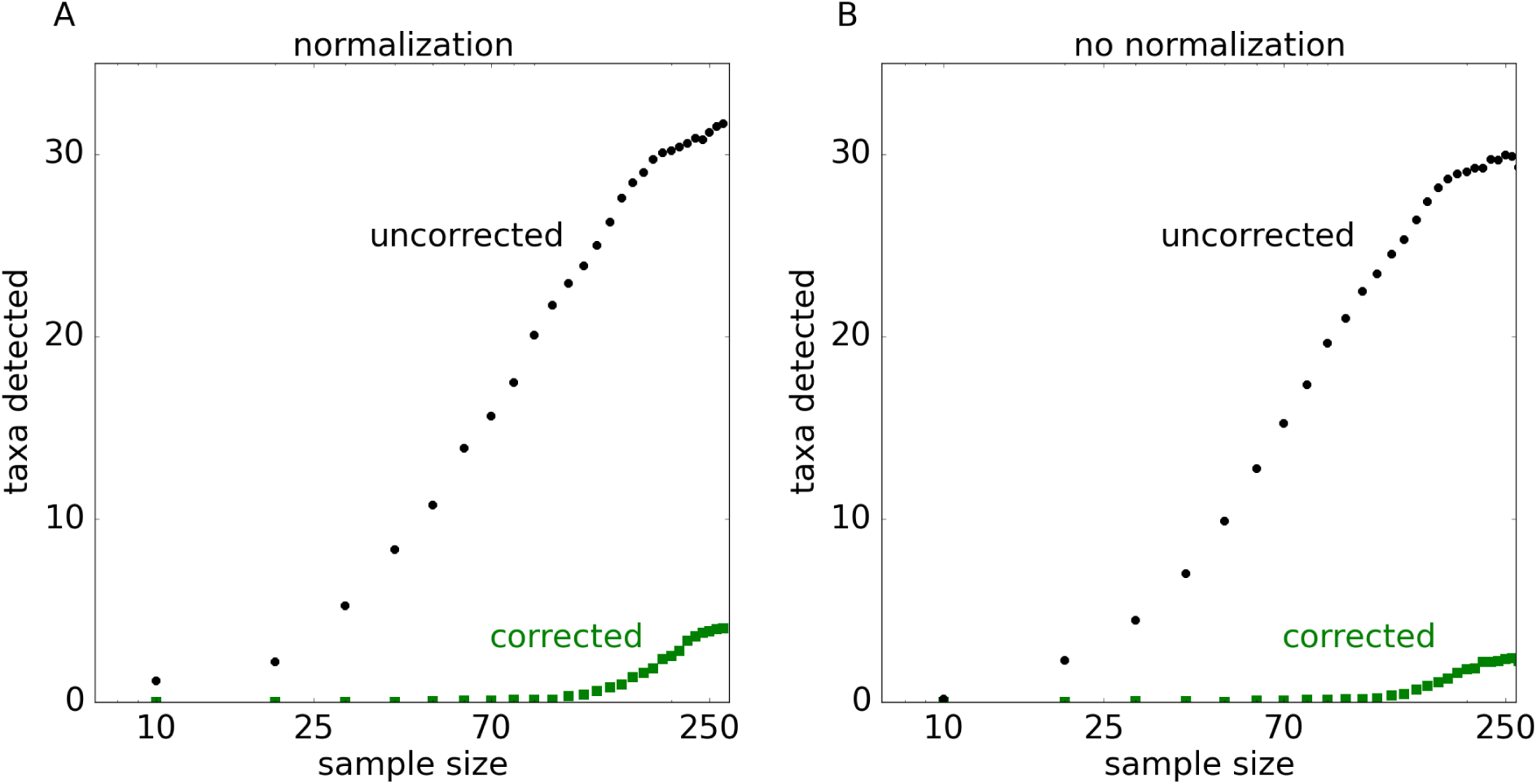
Compositional bias does not significantly affect DAA performance. **(A)** is the same as Fig. 2C of the main text. **(B)** is similar to (A), but with the analysis done on unnormalized counts, which do not add up to a constant number. The results of the two analyses are very similar suggesting that compositional bias does not create significant artifacts. In particular, the number of associations in (A) and in (B) grow at the same rate with the sample size. This would not be the case if the compositional bias was strong because spurious associations due to normalization would lead to a greater number of detected taxa. Thus, we conclude that interspecific interactions rather than compositional effects are the primary source of spurious associations.

**Figure S7.**
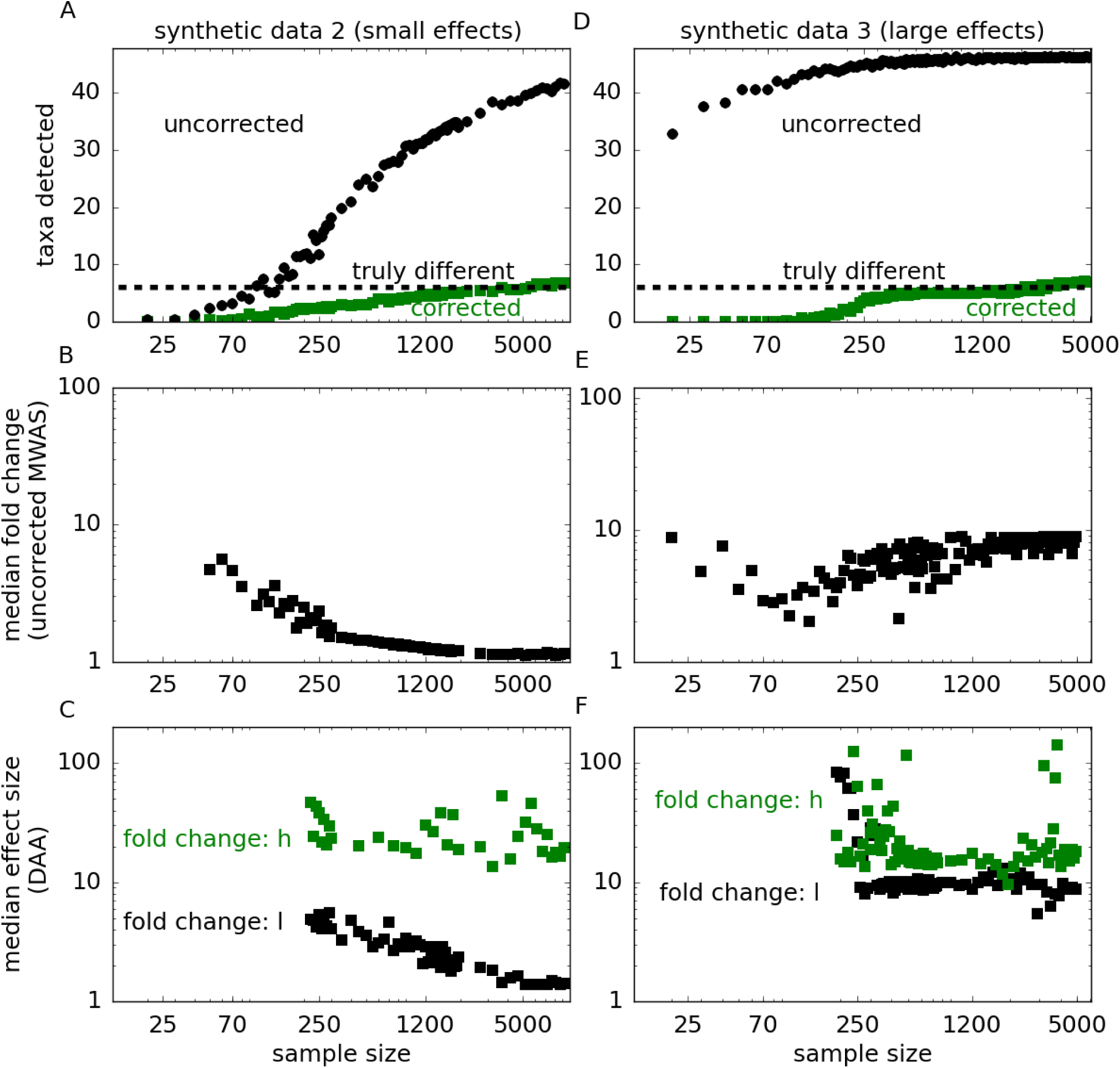
Spurious associations in synthetic data with small and large effect sizes. The same analysis as in Fig. 2AB of the main text, but for synthetic data with smaller (A, B, C) and larger (D, E, F) effect sizes. **(A)** and **(D)** show the number of associations detected by traditional MWAS and DAA. **(B)** and **(E)** show the median effect sizes (median fold change) for the taxa detected by conventional MWAS. **(C)** and **(E)** show the effect sizes in both h and l for the taxa detected by DAA. The effect size for h was quantified as the relative percent difference in host-field between cases and controls, while the l-effect size was computed as described in the main text. Overall the results are similar to those in Fig. 2. In addition, (A) and (B) show that DAA can recover all directly associated taxa given a large number of samples without any false positives. For sample sizes exceeding 5000, DAA starts to detect indirect associations due to compositional effects.

**Figure S8.**
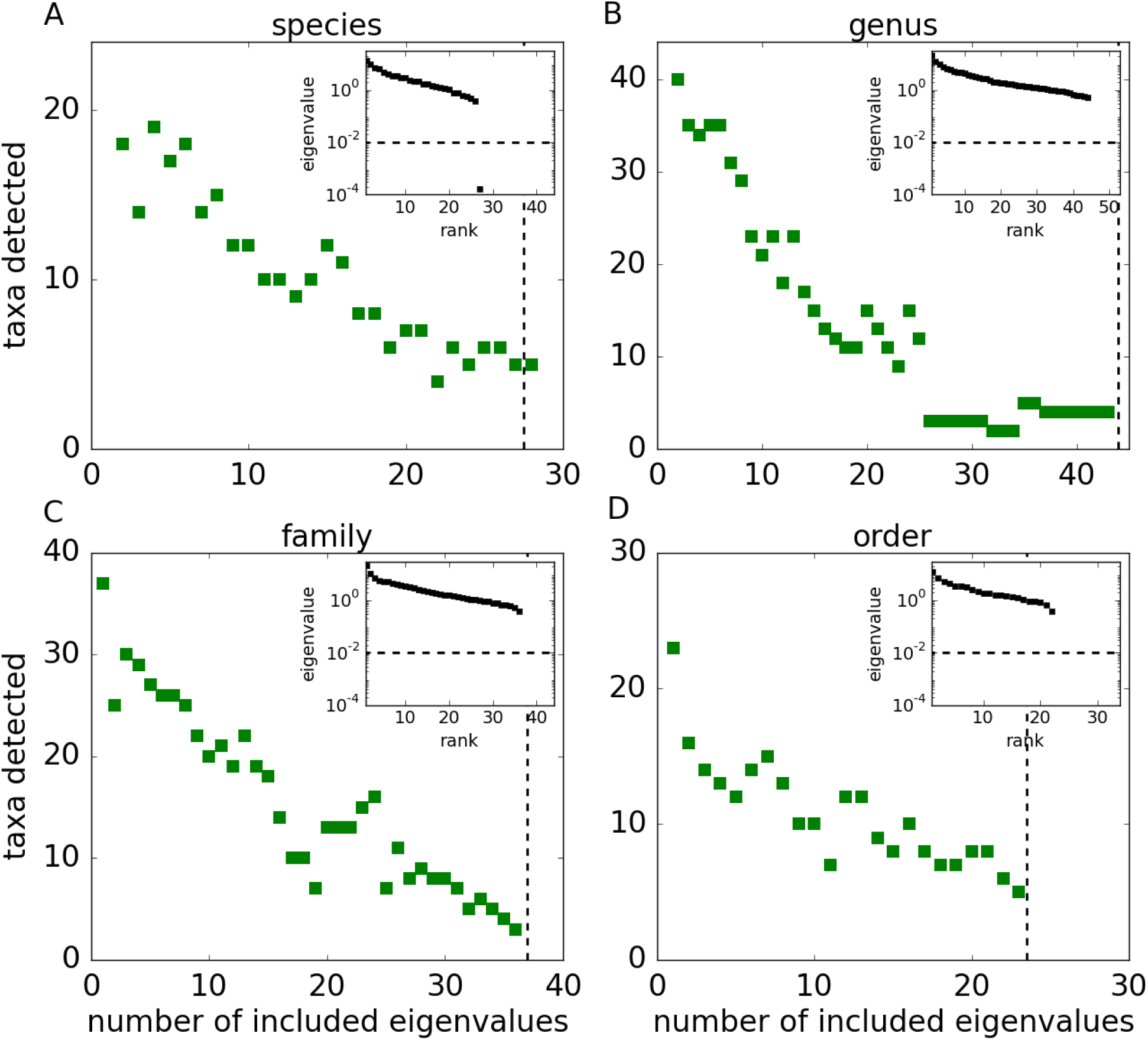
Sensitivity of DAA to eigenvalue threshold *λ*_min_. Large *λ*_min_ retains only a few eigenvalues and imposes an artificially strong correlation structure on the data. As a result, DAA detects a large number of associations because it cannot distinguish direct from indirect effects. The performance of DAA improves as more eigenvalues are included and reaches a plateau. The dashed lines show the number of eigenvalues included for *λ*_min_ = 0.01 used throughout our analysis. The insets show the eigenvalues of Λ in decreasing order.

**Table S1.** The list of genera used in the analysis. We included all genera that were present in more than 60% of either control or IBD subjects. The indices were chosen to hierarchically cluster the correlation matrix shown in Fig. 1b of the main text (index corresponds to the position of the genus on the x axis).

| index | genus name | index | genus name | index | genus name |
| --- | --- | --- | --- | --- | --- |
| 1 | *[Prevotella]* | 17 | *Corynebacterium* | 33 | *Fusobacterium* |
| 2 | *Prevotella* | 18 | *Pseudomonas* | 34 | *Bacteroides* |
| 3 | *Dialister* | 19 | *Acinetobacter* | 35 | *Anaerostipes* |
| 4 | *Phascolarctobacterium* | 20 | *Erwinia* | 36 | *Parabacteroides* |
| 5 | *Epulopiscium* | 21 | *Actinomyces* | 37 | *[Eubacterium]* |
| 6 | *Eggerthella* | 22 | *Streptococcus* | 38 | *Odoribacter* |
| 7 | *Clostridium* | 23 | *Granulicatella* | 39 | *Oscillospira* |
| 8 | *Akkermansia* | 24 | *Neisseria* | 40 | *Lachnospira* |
| 9 | *Bilophila* | 25 | *Rothia* | 41 | *Roseburia* |
| 10 | *Bifidobacterium* | 26 | *Eikenella* | 42 | *Faecalibacterium* |
| 11 | *Collinsella* | 27 | *Campylobacter* | 43 | *Dorea* |
| 12 | *Sutterella* | 28 | *Veillonella* | 44 | *[Ruminococcus]* |
| 13 | *Parvimonas* | 29 | *Actinobacillus* | 45 | *Ruminococcus* |
| 14 | *Porphyromonas* | 30 | *Aggregatibacter* | 46 | *Blautia* |
| 15 | *Turicibacter* | 31 | *Haemophilus* | 47 | *Coprococcus* |
| 16 | *Staphylococcus* | 32 | *Holdemania* |  |  |

**Table S2.** Genera modified in synthetic data. Taxa indices are the same as in Table S1. Effect size is the percent change in the value of *h*.

| taxon<br>index | effect size<br>data 1 (main text) | effect size<br>data 2 (small) | effect size<br>data 3 (large) |
| --- | --- | --- | --- |
| 1 | −18% | −17% | −44% |
| 11 | +24% | +14% | +129% |
| 19 | −36% | −12% | −72% |
| 27 | +17% | +16% | +67% |
| 33 | −13% | −14% | −28% |
| 45 | +18% | +13% | +112% |

**Table S3.** Direct associations identified by DAA across phylogenetic levels.

| taxon<br>name | direct effect,<br>$h_{CD}$ | direct effect,<br>$h_{ctrl}$ | difference,<br>$\Delta h/\vert h_{ctrl}\vert $ | p-value | q-value |
| --- | --- | --- | --- | --- | --- |
| **Order level** |  |  |  |  |  |
| *Burkholderiales* | -0.47 | -0.66 | +0.29 | 0.00013 | 0.0029 |
| *Turicibacterales* | -1.7 | -1.4 | -0.18 | 0.00031 | 0.0036 |
| *Pasteurellales* | -0.51 | -0.69 | +0.26 | 0.00068 | 0.0052 |
| *Campylobacterales* | -1.6 | -1.8 | +0.1 | 0.00696 | 0.04 |
| *Erysipelotrichales* | -2.5 | -2.3 | -0.083 | 0.0095 | 0.044 |
| **Family level** |  |  |  |  |  |
| *Alcaligenaceae* | -0.68 | -0.86 | +0.21 | 0.00027 | 0.01 |
| *Clostridiaceae* | -1.2 | -0.99 | -0.18 | 0.0026 | 0.049 |
| *Pasteurellaceae* | -0.31 | -0.47 | +0.35 | 0.0033 | 0.049 |
| **Genus level** |  |  |  |  |  |
| *Roseburia* | -1.2 | -0.86 | -0.35 | 0.000098 | 0.0046 |
| *Sutterella* | -0.63 | -0.80 | +0.22 | 0.00043 | 0.01 |
| *Oscillospira* | -2.4 | -2.6 | +0.097 | 0.0015 | 0.023 |
| *Turicibacter* | +0.46 | +0.69 | -0.34 | 0.003 | 0.035 |
| **Species level** |  |  |  |  |  |
| *B.adolescentis* | -0.23 | +0.073 | -4.12 | 0.00013 | 0.0037 |
| *E.dolichum* | -0.51 | -0.31 | -0.65 | 0.0028 | 0.039 |
| *F.prausnitzii* | -0.97 | -0.81 | -0.20 | 0.0042 | 0.039 |
| *A.segnis* | -0.072 | -0.25 | +0.71 | 0.0056 | 0.04 |
| *B.producta* | -0.75 | -0.54 | -0.38 | 0.0064 | 0.04 |

**Table S4.** Comparison between changes in *h* and in *l* for the taxa identified by DAA.

| taxon<br>name | abundance<br>$l_{\text{CD}}/l_{\text{ctrl}}$ | direct effect<br>$\Delta h/\vert h_{\text{ctrl}}\vert $ | q-value, $l$ | q-value, $h$ |
| --- | --- | --- | --- | --- |
| **Order level** |  |  |  |  |
| *Burkholderiales* | +1.6 | +0.29 | 0.04 | 0.0029 |
| *Turicibacterales* | +0.45 | −0.18 | 0.00002 | 0.0036 |
| *Pasteurellales* | +4.2 | +0.26 | 0 | 0.0052 |
| *Campylobacterales* | +2.1 | +0.1 | 0.000001 | 0.04 |
| *Erysipelotrichales* | +0.34 | −0.083 | 0 | 0.044 |
| **Family level** |  |  |  |  |
| *Alcaligenaceae* | +1.7 | +0.21 | 0.03 | 0.01 |
| *Clostridiaceae* | +0.25 | −0.18 | 0 | 0.049 |
| *Pasteurellaceae* | +4.2 | +0.35 | 0 | 0.049 |
| **Genus level** |  |  |  |  |
| *Roseburia* | +0.21 | −0.35 | 0 | 0.0046 |
| *Sutterella* | +2.0 | +0.22 | 0.004 | 0.01 |
| *Oscillospira* | +0.84 | +0.097 | 0.33 | 0.023 |
| *Turicibacter* | +0.50 | −0.34 | 0.0004 | 0.035 |
| **Species level** |  |  |  |  |
| *B.adolescentis* | +0.43 | −4.12 | 0.00004 | 0.0037 |
| *E.dolichum* | +0.43 | −0.65 | 0.00004 | 0.039 |
| *F.prausnitzii* | +0.41 | −0.20 | 0.000003 | 0.039 |
| *A.segnis* | +2.8 | +0.71 | 0 | 0.04 |
| *B.producta* | +0.67 | −0.38 | 0.03 | 0.04 |

**Table S5.**
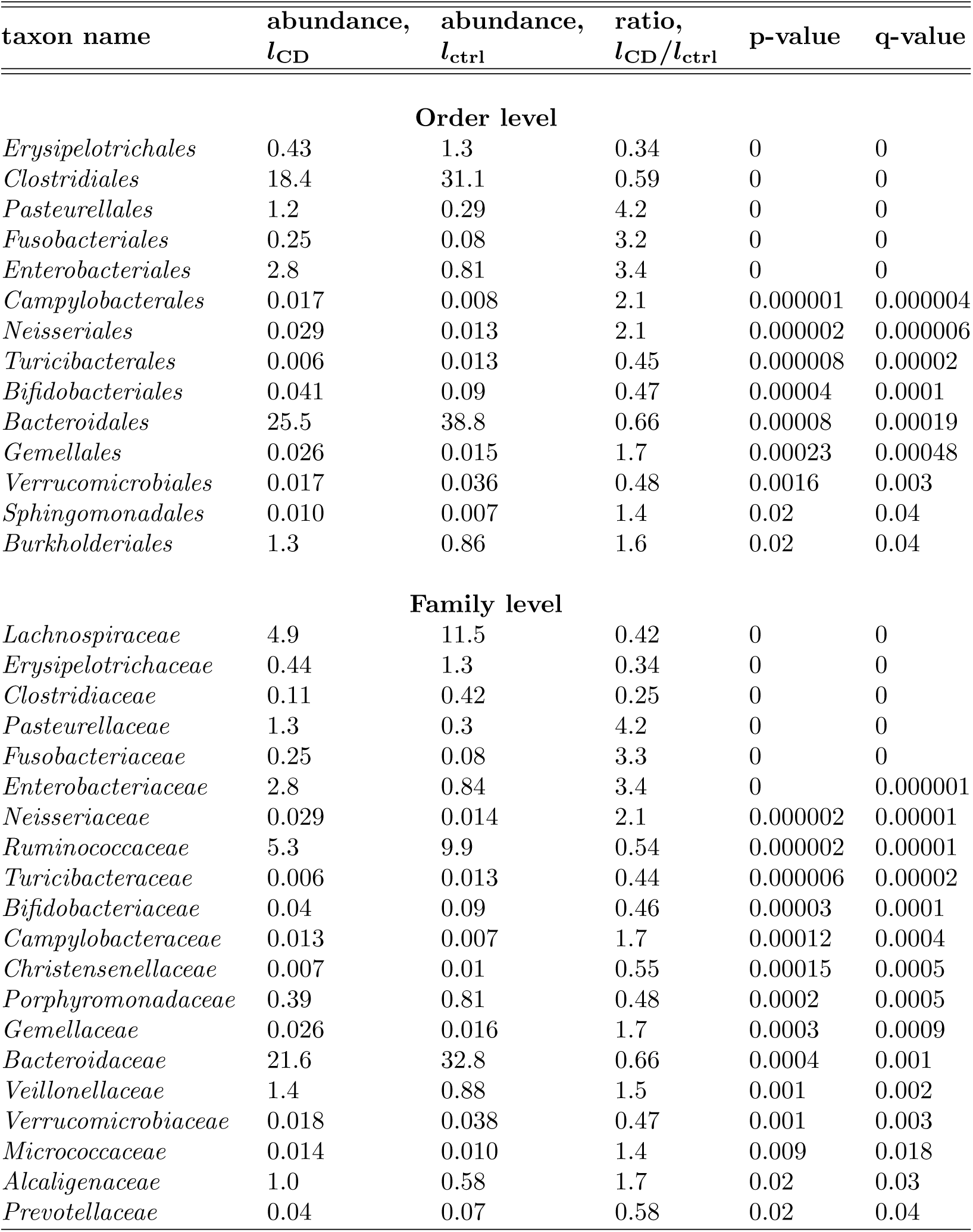
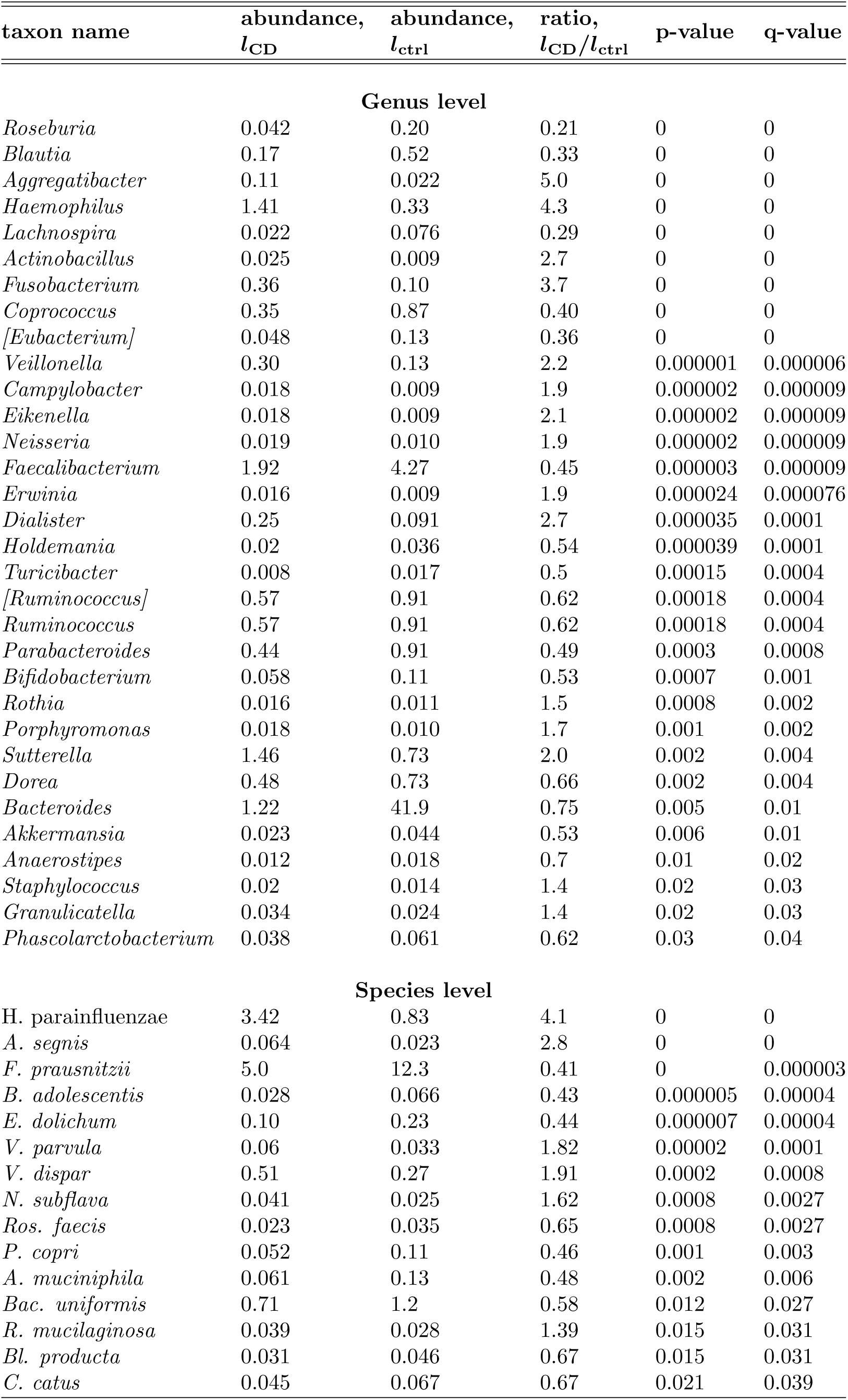
Indirect associations identified by uncorrected abundance analysis across phylogenetic levels.

**Table S6.** A summary of interaction strengths and log-abundance correlation coefficients for the core IBD network shown in Fig. 3 of the main text. Statistical significance was estimated by a permutation test. Specifically, we independently permuted the abundance of each taxa across samples and then computed the correlation and interaction matrices on the permuted data to generate the probability distribution for the null hypothesis of no interaction.

| interacting taxa | correlation<br>strength, $C_{ij}$ | interaction<br>strength, $J_{ij}$ | q-value,<br>correlation | q-value,<br>interaction |
| --- | --- | --- | --- | --- |
| *A.segnis-B.producta* | +0.16 | +0.14 | 0.0011 | 0.0041 |
| *A.segnis-Oscillospira* | -0.16 | -0.17 | 0.0014 | 0.0011 |
| *A.segnis-Roseburia* | -0.15 | -0.19 | 0.0034 | 0.0006 |
| *A.segnis-Sutterella* | -0.015 | +0.046 | 0.80 | 0.41 |
| *A.segnis-Turicibacter* | +0.18 | +0.12 | 0 | 0.021 |
| *B.adolescentis-A.segnis* | +0.19 | +0.19 | 0 | 0.0006 |
| *B.adolescentis-B.producta* | +0.26 | +0.16 | 0 | 0.0019 |
| *B.adolescentis-Oscillospira* | +0.069 | -0.067 | 0.17 | 0.24 |
| *B.adolescentis-Roseburia* | +0.25 | +0.24 | 0 | 0 |
| *B.adolescentis-Sutterella* | +0.036 | +0.055 | 0.50 | 0.34 |
| *B.adolescentis-Turicibacter* | +0.40 | +0.46 | 0 | 0 |
| *B.producta-Oscillospira* | +0.10 | +0.04 | 0.044 | 0.47 |
| *B.producta-Roseburia* | +0.100 | +0.0063 | 0.047 | 0.92 |
| *B.producta-Sutterella* | +0.0012 | +0.092 | 0.98 | 0.091 |
| *B.producta-Turicibacter* | +0.31 | +0.23 | 0 | 0 |
| *E.dolichum-A.segnis* | -0.0063 | -0.027 | 0.92 | 0.66 |
| *E.dolichum-B.adolescentis* | +0.19 | +0.051 | 0.0002 | 0.35 |
| *E.dolichum-B.producta* | +0.40 | +0.46 | 0 | 0 |
| *E.dolichum-F.prausnitzii* | +0.075 | +0.0087 | 0.13 | 0.92 |
| *E.dolichum-Oscillospira* | +0.27 | +0.29 | 0 | 0 |
| *E.dolichum-Roseburia* | +0.25 | +0.21 | 0 | 0 |
| *E.dolichum-Sutterella* | -0.080 | -0.19 | 0.11 | 0 |
| *E.dolichum-Turicibacter* | +0.20 | +0.057 | 0 | 0.33 |
| *F.prausnitzii-A.segnis* | -0.086 | +0.0064 | 0.086 | 0.92 |
| *F.prausnitzii-B.adolescentis* | +0.15 | +0.20 | 0.0021 | 0 |
| *F.prausnitzii-B.producta* | -0.065 | -0.15 | 0.19 | 0.0032 |
| *F.prausnitzii-Oscillospira* | +0.32 | +0.29 | 0 | 0 |
| *F.prausnitzii-Roseburia* | +0.35 | +0.35 | 0 | 0 |
| *F.prausnitzii-Sutterella* | +0.25 | +0.204 | 0 | 0.0006 |
| *F.prausnitzii-Turicibacter* | -0.095 | -0.18 | 0.053 | 0.0003 |
| *Roseburia-Oscillospira* | +0.29 | +0.16 | 0 | 0.0034 |
| *Roseburia-Sutterella* | +0.099 | +0.019 | 0.05 | 0.76 |
| *Roseburia-Turicibacter* | +0.099 | +0.053 | 0.05 | 0.34 |
| *Sutterella-Oscillospira* | +0.23 | +0.24 | 0 | 0 |
| *Turicibacter-Oscillospira* | +0.036 | +0.076 | 0.50 | 0.18 |
| *Turicibacter-Sutterella* | -0.12 | -0.15 | 0.012 | 0.0026 |

